# Indenting multi-cellular spheroids with various cantilever tip geometry

**DOI:** 10.1101/2025.07.05.663257

**Authors:** Kajangi Gnanachandran, Ewelina Lorenc, Alessandro Podestà, Małgorzata Lekka

## Abstract

Spheroids are of great interest in the study of cancer as they can partially mimic the tumour microenvironment, thus allowing to investigate several aspects of cell – microenvironment interactions in healthy and diseased conditions, including those pertaining to mechanobiology. Atomic Force Microscopy (AFM) is a versatile tool for studying biological samples and their mechanobiological properties. In AFM, the tip shape and dimensions determine the contact geometry between the tip and the sample and the length scales at which the mechanical properties are probed. Given the complex multiscale structure of spheroids, the choice of tip geometry and size would allow, in principle, to dissect the mechanical response of the overall system into the contributions of the constituents, from the single cell level to the cellular aggregate. In this work, we studied the mechanical properties of spheroids derived from four cell lines (A549, NHLF, HT-29, CCD-18Co). Our studies revealed that using different contact geometries in the fitting procedure results in significantly different Young’s modulus values, highlighting the multiscale response of these complex cellular systems and the importance of a precise experiment design and choice of the AFM probe for the nano-mechanical measurements. We observed that the location of F-actin filaments is correlated to the rigidity of the spheroids.

## 1. Introduction

In recent years, multi-cellular spheroids have been widely used as a good in vitro model in various types of research, especially in drug efficacy and toxicity trials, but also in the investigation of organ development and congenital diseases, tissue engineering, and 3D bioprinting [1–3]. Spheroids are now the most desirable 3D model for creating uniform, repeatable multi-cellular structures and a potential starting point for building big tissues and intricate organs [4]. The spheroids are of big interest in the study of cancer as they can partially mimic the tumour microenvironment allowing a better recapitulation of the patient tumour, which represents a promising challenge to improve the success rates in anticancer drug development and study of cell-extracellular matrix (ECM) interaction, cancer biology, including cancer initiation, invasion, and metastatic processes [4,5].

Cancer cells are known to be subjected to variable stresses during cancer progression due to the fast-dividing process in a restricted space [6]. Likewise, several mechanical events take place during tumour progression. For instance, there is an overproduction of collagen by stromal myofibroblasts that causes stiffening in the matrix [7]; the tumour mass expansion induces compression [8], and because of the poor lymphatic drainage, there is a rise of the interstitial pressure [9]. Several studies have shown how spheroids are a potentially powerful model to study the influence of mechanical stress on tumour growth [10–12].

Since it has been demonstrated that the mechanical properties of cancerous cells and the organisation of their cytoskeletal network are altered compared to normal cells [13,14], the mechanobiology of cells has reached an important role in cancer research along with the traditional genetic and biochemical studies. There is a broad selection of tools for studying the mechanical properties of cells, including Atomic Force Microscopy (AFM), micropipette aspiration, optical and magnetic tweezers, and different microfluidic approaches [15]. AFM is a powerful tool that, thanks to its versatility, has been used to study a variety of biological samples and different mechanobiological aspects [16], by characterizing their viscoelastic [17–19] and adhesive properties [20]. However, only recently, the mechanical properties of spheroids have become of interest in AFM studies due to their complex and multilayer structure [21–23].

The key elements of the AFM are the cantilever and the probing tip [24]. Since different tip geometries are appropriate for various uses and sample types, the study objectives and the sample properties will ultimately define which tip geometries should be used. The tip interacts directly with the sample, and its shape determines the contact geometry between the tip and the sample. Contact mechanics relate the load force F and the indentation δ [25] [26]. The obtained equations take into account the geometry of the indenter, i.e., the probing tip shape (Table 1) [27].

**Table 1.**
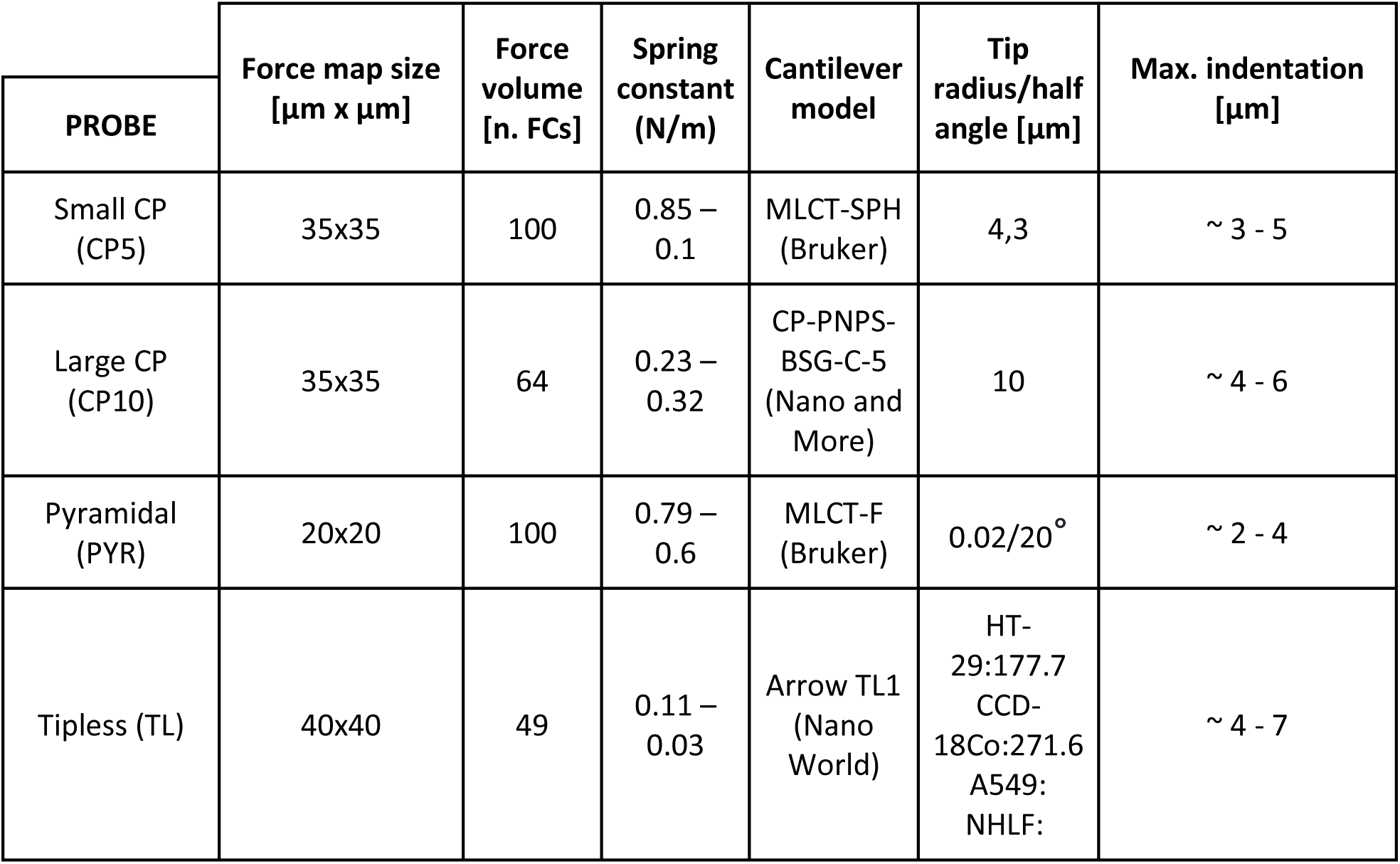
AFM tips used for experiments and measurement parameters.

| PROBE | Force map size<br>[$\mu\text{m} \times \mu\text{m}$] | Force<br>volume<br>[n. FCs] | Spring<br>constant<br>(N/m) | Cantilever<br>model | Tip<br>radius/half<br>angle [$\mu\text{m}$] | Max. indentation<br>[$\mu\text{m}$] |
| --- | --- | --- | --- | --- | --- | --- |
| Small CP<br>(CP5) | 35x35 | 100 | 0.85 –<br>0.1 | MLCT-SPH<br>(Bruker) | 4,3 | ~ 3 - 5 |
| Large CP<br>(CP10) | 35x35 | 64 | 0.23 –<br>0.32 | CP-PNPS-BSG-C-5<br>(Nano and<br>More) | 10 | ~ 4 - 6 |
| Pyramidal<br>(PYR) | 20x20 | 100 | 0.79 –<br>0.6 | MLCT-F<br>(Bruker) | 0.02/20° | ~ 2 - 4 |
| Tipless (TL) | 40x40 | 49 | 0.11 –<br>0.03 | Arrow TL1<br>(Nano<br>World) | HT-29:177.7<br>CCD-18Co:271.6<br>A549:<br>NHLF: | ~ 4 - 7 |

Pyramidal tips are the most suitable choice for experiments that aim to obtain high-resolution images of scanned surfaces or to create force maps [28]. The use of pyramidal tips allows for deep indentation studies and for imaging small features of the sample. Unfortunately, in the case of studies on biological samples, sharp tips are a source of high stress [29]. Colloidal probes (CP), with radii between 2 and 20 µm, represent an alternative to study the mechanical properties of biological samples [24,30,31]. First of all, CPs do not cause high stress even for high indentations, which can, therefore be higher than in the case of pyramidal tips. The cost of using CP is lower lateral resolution; however, this issue can be overcome by using CP of smaller diameter [29–31]. The increased contact area between the sphere and sample allows for detecting - weak interaction forces, which is another characteristic of CPs. In general, CPs make it easier to interpret data regarding theoretical models [30,31]. To perform measurements with the constant and defined area, cylindrical tips can be used [32]. As they provide a linear force-indentation relationship, cylindrical tips find multiple applications in studies of cell adhesion and nonlinear response of cells [26].

Several studies have already reported the effects of different types of cantilevers on the elastic properties of single cells. These studies have demonstrated that the measured value of Young’s modulus (YM) of elasticity depends on the cantilever type and on the used contact mechanics models that rely on several approximations [29,33,34]. These are among the main reasons for the variability of the published results [35]. Most of them were conducted on 2D cell cultures and there are no data showing analogous effects of the cantilever geometry on the mechanical properties of 3D systems like spheroids. To fill the gap, we performed AFM-based elasticity measurements on spheroids formed from four distinct cell lines characterised by large variations in the organisation of their actin cytoskeleton, which is either well- or poorly differentiated. To get the mechanical response from the spheroids at different scales, we used four types of cantilevers equipped with tips of different sizes and geometry. Our results indicate that regardless of the tip used, the spheroids possessing similar actin cytoskeleton organisation do not display any mechanical differences, indicating that even at the spheroid level, the mechanical properties of cells are related to their cytoskeletal organisation.

## 2. Materials and Methods

### 2.1. Cell culture

In this research, four different cell lines were used: A549 (human lung carcinoma, Lonza) and NHLF (human lung fibroblasts, ATCC), HT-29 (human colon adenocarcinoma, a kind gift from the Chair of Medical Biochemistry, Collegium Medicum Jagiellonian University, Cracow, Poland), and CCD18-Co (human colon fibroblasts, a kind gift from the Department of Virology and Immunology, University of Maria Sklodowska-Curie, Lublin, Poland). NHLF cells were cultured in Fibroblast basal medium (FBM) + Fibrobalst growth medium -2 (FGM-2), while A549 cells were cultured in F12K medium. Both culture media were supplemented with 10 % Fetal bovine serum (FBS). HT-29 and CCD18-Co cells were cultured in RPMI 1640 and Advanced DMEM medium, supplemented with 5% FBS, 1% penicillin/streptomycin, and 1% amphotericin. The lung cell lines were grown in culture flasks (Sarstedt) in an incubator (Nuaire) at 37° C with 5% CO_2_. HT-29 and CCD18-Co cells were grown in the same conditions in the incubator (Galaxy S, RS biotech). The passaging was carried out when the cells reached 80-90% of the confluency level. A 0.25% trypsin-EDTA solution was used for all cell lines to detach them from the culture flask surface. If not stated otherwise, all bioreagents and materials came from Sigma Aldrich.

### 2.2. Confocal microscopy

Confocal images of actin filaments inside spheroids were visualised using a confocal microscope, accessible in the Laboratory *of in vivo* and *in vitro* imaging (Maj Institute of Pharmacology, Polish Academy of Sciences, Cracow, Poland). The samples were stained using the following protocol. Spheroids formed by each cell line were collected in a 1.5 ml Eppendorf and fixed with 3.7% paraformaldehyde for 1 hour. The samples were washed with phosphate bufferes saline (PBS) for 2 minutes three times, treated with 1% cold Triton X-100 overnight at 4°C, and washed again with PBS. Then, 1% cold Bovine Serum Albumin solution (BSA, Sigma-Aldrich) was added for 3 hours at 4°C, and after removing BSA, the spheroids were washed with PBS. Next, the spheroids were incubated overnight at 4°C with phalloidin conjugated with Alexa Fluor 488 (1:20 in PBS). The next day, the dye for actin filaments was removed, and Hoechst 33342 (Invitrogen, dissolved in a PBS buffer (1:5)) staining the cell nuclei was added. Finally, the samples were washed with PBS and moved to 18-well glass-bottom slides (Ibidi) with an anti-shading solution (Thermofisher) having the same refractive index (1.52) as the oil used for the objective of 63x magnification. Images were recorded using a Leica TCS SP8 WLL confocal microscope equipped with new-generation HyD detectors. Fluorescent dyes were excited by diode lasers at 405 nm (Hoechst) and white light laser with emission wavelength set at 499 nm (Alexa Fluor 488). Images were registered using an oil immersion 63x objective lens (HC PL APO CS2 NA 1.40).

### 2.3. Preparation and fixation of spheroids for AFM measurements

The spheroids were cultured using a low attachment method using U-bottom 96-well plates (Thermo Fisher). To obtain spheroids of size around 350-400 µm in diameter, adequate numbers of cells/well were chosen. These were 1500 cells/well for A549 lung cancer cells, 1800 cells/well for NHLF lung fibroblasts, 1000 cells/well for HT-29 colon cancer cells, and 10000/well for CCD-18Co colon fibroblasts. After 3 days of culture, the diameter of spheroids was determined based on bright field images using ImageJ software. To keep the spheroids immobile during the indentation, the Petri dishes were coated with 0.01% poly-lysine and 5% glutaraldehyde [35] with an addition support of a Micromesh array (Microsurfaces Pty Ltd, Australia). This was fixed on the Petri dish following the producers’ instructions, and the dish was then filled with adequate cell culture medium without phenol red. Spheroids were transferred from a multi-well culture plate to the array, placing each spheroid in a well, which makes measurements easier. Placing all the spheroids under AFM steup (outside of culturing conditions) for the time of measurements (several hours) could affect spheroids properties and quality of measurements. Considering this, only several spheroids were collected at time and after measuring 3-4 of them, new spheroids were collected to continue measurements.

### 2.4. AFM tip geometries

Four cantilever types with different tip geometries were used to study the mechanical properties of spheroids (**Table 1**). Two types of triangular cantilevers with spherical tips (CP5 and CP10, with radii of 5 and 10 µm, respectively), triangular cantilevers (MLCT-F) with a four-sided pyramidal tip (open-angle of 20° and radius 20-40 nm), and rectangular tipless cantilevers (Arrow TL1). For the MLCT cantilevers, the tip height varied between 3 µm and 8 µm with a mean of 5.5 µm (according to manufacturer data). Prior to each experiment, the spring constants were calibrated with the thermal noise methods [36]. The radii of the CPs were either pre-calibrated by the manufacturer or calibrated using the procedures described in Indrieri et al. [37]. The radii of the spheroids were determined using an optical microscope.

In Table 1 the details about the various tips and the maximum indentation levels are reported.

### 2.5. AFM measurements

For spheroids formed by human colon HT-29 and CCD-18Co cells, we used a Bioscope Catalyst AFM (Bruker). A Nanowizard IV AFM (Bruker – JPK Instruments) was used to measure the spheroids formed by the A549 and NHLF human lung cells. The instruments were mounted on top of an inverted optical microscope (Olympus IX71). To isolate the sample from the ground and ambient noise, the microscope was placed on an active anti-vibration base (DVIA-T45, Daeil Systems) inside an acoustic enclosure (Schaefer).

Spheroids were measured at room temperature (around 24°C) in their respective culture medium without phenol red. The medium was changed several times to avoid heating it up from the microscope lamp. In all measurements, the approach speed was set to 10 µm/s, and the maximum load force was kept at 15 nN. Two different indentation levels with the pyramidal tip were obtained by adjusting the load force to 5 nN and 15 nN, respectively, for low and high indentation. For each set of measurements, 10 spheroids were measured, and 3-6 force maps were collected per spheroid. A suitable scan area was chosen according to the contact area of the different cantilevers, as indicated in **Table 1**.

### 2.6. Young’s modulus determination and statistics

The data obtained from spheroids formed by A549 and NHLF cells were processed with the software provided by JPK. The data obtained for spheroids derived from the other colon cell types were analysed by using custom MATLAB scripts following the routine described in Puricelli et al. [31]. In both cases, the rescaled force-indentation curves were fitted using the appropriate contact mechanics models (**Figure 1**) to obtain the values of Young’s modulus [38]. The Hertz model for the paraboloidal indenter was used as an approximation of the model for the spherical indenter [26].

**Figure 1.**
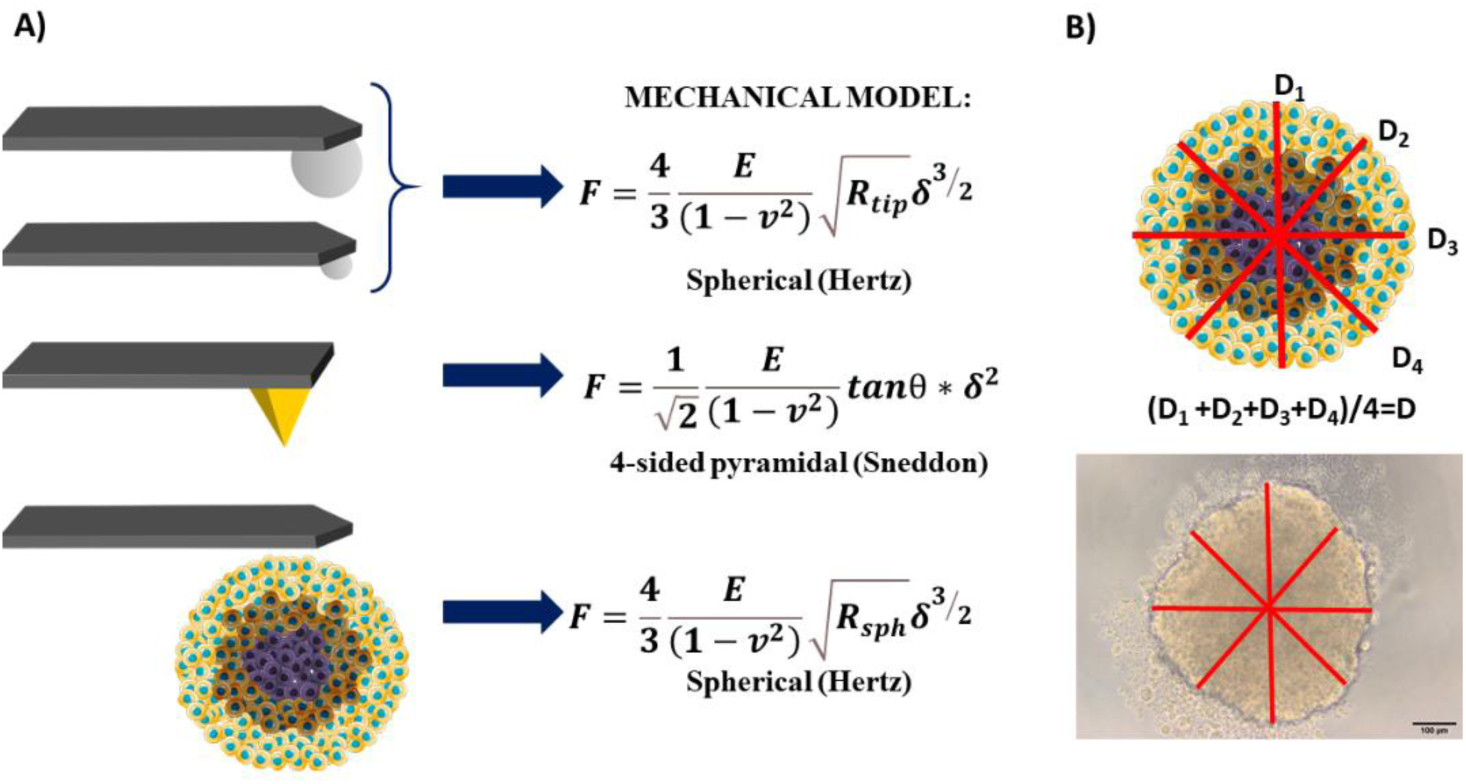
A) Summary of the selected contact mechanics models for the AFM measurements on spheroids performed with various tip geometries (*R_tip_* - radius of the beads; *R_sph_* - average radius of spheroids). B) Determination of the spheroid diameter.

In the case of measurements with tipless cantilevers, the Hertz model for spherical probes was applied, assuming the indentation of a soft spherical body (the spheroid) by a flat and stiff plate (the tipless cantilever). As the radius of the spherical body, we took the average radius of the spheroids (approximately 300 μm). Additionally, we performed trial data analysis using two additional models. The first is the flat punch model (1) [29], assuming the cantilever width *a* as the diameter of the cylinder (therefore the punch radius is a’=a/2):

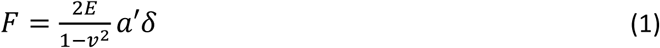

The other model is the Hertz model for colloidal probes, where we assumed the spheroid as locally flat and the tipless cantilever as an effective spherical indenter with diameter equal to the cantilever width. The statistical significance was calculated using an ordinary ANOVA test with Turkey’s multiple comparison test.

## 3. Results and discussions

### 3.1. Organization of F-actin inside the spheroids formed by human lung and colon cells

The mechanical properties of cells are predominantly related to the organization of the actin cytoskeleton, not only at the single-cell level [39] but also at the spheroid level [18]. Therefore, in this study, we first performed confocal imaging of the spheroids formed by the healthy lung fibroblasts (NHLF), colon fibroblasts (CCD18-Co), the small-lung cancer cells (A549), and colon cancer cells (HT-29), which were stained with Hoechst 33342 and Alexa Fluor 488 to observe the cell nucleus and the organization of actin filaments inside spheroids, respectively. In **Figure 2**, the confocal images of all the spheroids are shown.

**Figure 2:**
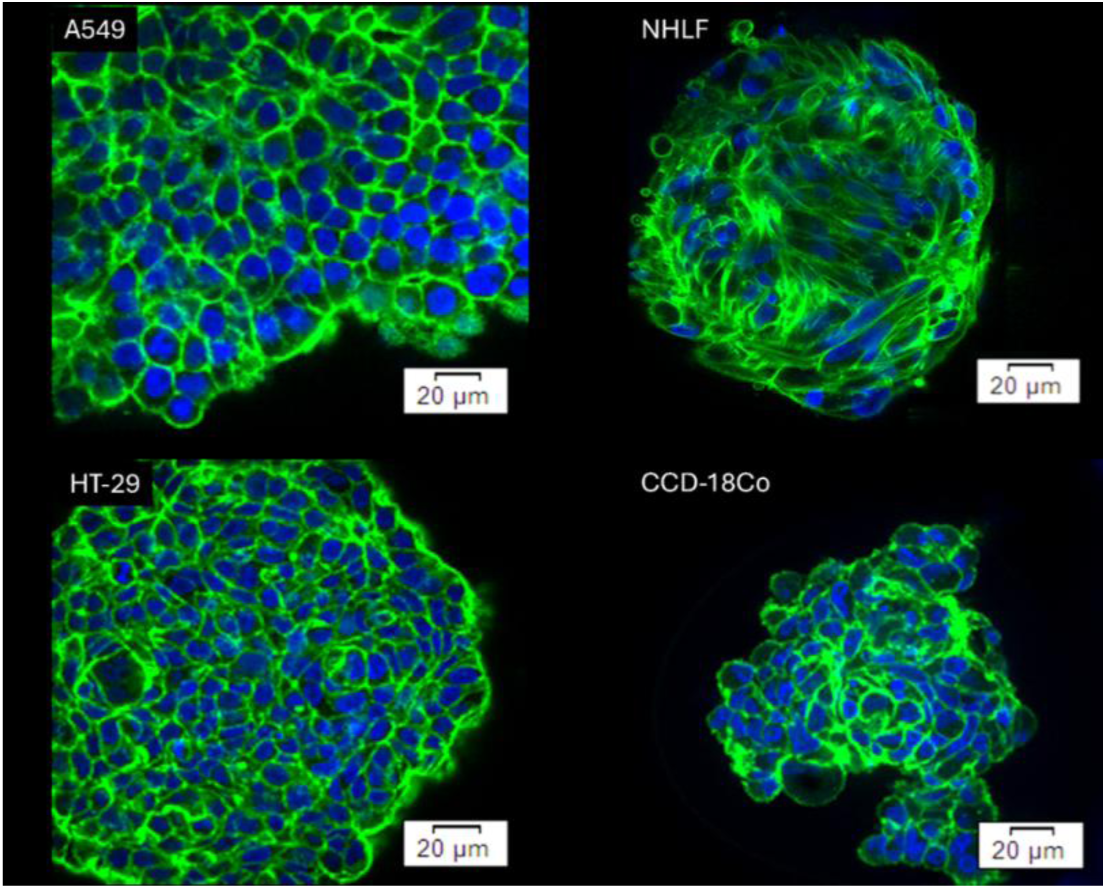
Confocal images of spheroids, formed by (A) lung cancer cells (A549 cell line), (B) healthy lung fibroblasts (NHLF cell line), (C) colon cancer (HT-29 cell line), and (D) colon fibroblasts (CCD-18Co cell line) [one slice from the Z-stack is shown]. Blue: nucleus (Hoechst 33342), Green: actin filaments (phalloidin Alexa Fluor 488).

In the spheroids formed by lung A549 cells, actin filaments are organized around the cells. Compared to A549 spheroids, those formed by the healthy lung fibroblasts (NHLF) present thick actin bundles distributed around and over the cells. The confocal images show a dissimilarity in the shapes of individual cells, i.e., more rounded A549 cells and more elongated NHLF fibroblasts. The spheroids formed by colon epithelial (HT-29 cells) and colon fibroblasts (CCD-18Co cells) display an actin filaments organization similar to those present in the spheroids formed by A549 cells, with actin filaments mainly distributed around the spherical cells.

To sum up, colon spheroids mainly consist of round cells with F-actin located around the cells, while lung spheroids are composed of spindle-like fibroblasts with actin filaments spanning over a whole cell or round, epithelial cells with F-actin organisation similar to those observed in colon spheroids.

### 3.2. Mechanical properties of spheroids formed by human lung and colon cells

The influence of the cantilever tip geometry on AFM nano-mechanical measurements has already been investigated on individual cells cultured on flat 2D surfaces, as shown in one of the most recent studies on NIH/3T3 fibroblasts [40]. Here, we take a step forward by performing a similar study at the 3D level. Therefore, this will allow us to understand the mechanical properties in a biological model, which is closer to the actual tumour microenvironment. The mechanical results for the spheroids formed by A549 and NHLF lung cells as well as by HT-29 and CCD18Co colon cells are reported in **Figure 3**.

**Figure 3:**
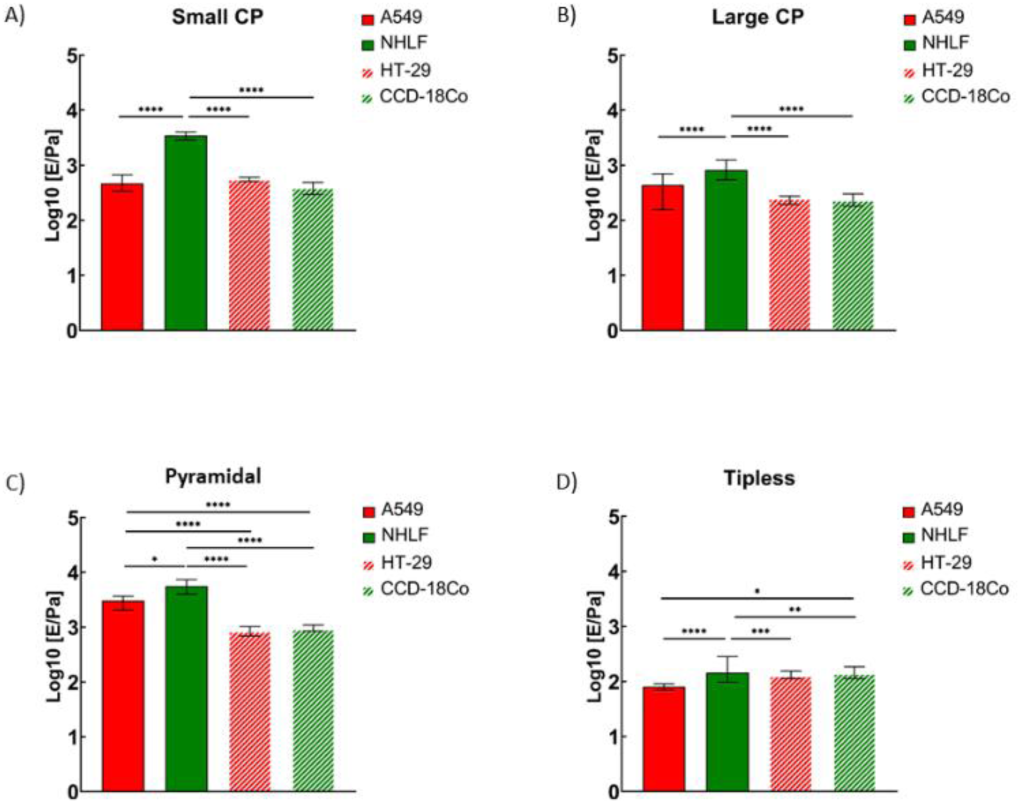
Young’s modulus obtained for spheroids formed from lung (A549 and NHLF) and colon (HT-29 and CCD18-Co) cells. Each bar represents the mean value ± standard error of the mean. Results obtained with A) Cantilevers with spherical probes of 4.3 µm (CP5) with indentations of 2-3 µm; B). Cantilevers with 10 µm (CP10) spherical probes with 4-6 µm indentations. C) MLCT indentation (2-4 µm cantilevers at); D) Tipless cantilevers (TL) with indentation 4-6 µm *(**** p <* 0.0001, *ns* - not statistically significant).

### 3.3. Colloidal probes

The mechanical properties of lung and colon spheroids were first measured using two distinct colloidal probes (CP5 and CP10) to probe the mechanical response of spheroids on different scales **(Figure 3A-B**).

In our measurements, the indentation achieved allowed us to study the outermost layers of the spheroids, i.e., the first few layers of cells and ECM produced by the cells.

The YM, obtained from spheroids formed from A459 lung cancer cells, was independent of the spherical probes used in the measurements. Its value remained similar, i.e., 0.50 ± 0.03 kPa (n = 39 maps) versus 0.45 ± 0.06 kPa (n = 31 maps), for spherical probes of 5 µm and 10 µm radius, respectively. A different situation is observed for spheroids formed from NHLF lung fibroblasts. Here, YM was 3.60 ± 0.27 kPa (n = 30 maps) versus 0.95 ± 0.09 kPa (n = 33 maps) for spherical probes of 5 µm and 10 µm radius, respectively.

Spheroids derived from colon cell lines were softer than those from lung cell lines. YM values measured with a spherical probe of radius 5 µm were 0.58 ± 0.03 kPa (n = 61) for HT-29 spheroids and 0.49 ± 0.08 kPa (n = 57) for CCD-18Co spheroids. With a larger sphere, obtained values for HT-29 and CCD-18Co spheroids were 0.31 ± 0.03 kPa (n = 59) and 0.31 ± 0.05 kPa (n = 50), respectively. Both results indicate that in the case of colon cells, there is no difference in elastic properties between spheroids derived from cancerous cells and fibroblasts.

### 3.4. Pyramidal cantilevers

The mechanical properties of spheroids were then measured with pyramidal tips (**Figure 3C**). YM of lung cancer cells and fibroblasts was measured at two different indentation depths (low indentation = 1 µm; high indentation: 2-3 µm), which were obtained by adjusting the load force to 5 nN and 15 nN, respectively. They were smaller than the values obtained with the colloidal probes. Two different indentation ranges for HT-29 and CCD-18Co were extracted from two fitting ranges: 10%-30% and 30%-90% of linearized force curves. Since we did not observe significant differences in YM in those two indentation ranges (**Supplementary Figure S1**) we only show the high indentation results.

**Figure 3C** shows that for each cell line, YM remained the same, regardless of the indentation depths considered. It suggests that the probed spheroid layer remained mechanically homogenous. On the other hand, the lack of differences can be explained by the fact that the thickness of the probed layer is not very high, as the height of the pyramidal tip limits it. Moreover, the spheroid radius is much larger than the maximum indentation allowed by the standard AFM setup, therefore, our measurement has a surface sensitivity rather than bulk sensitivity; this condition is clearly different from the typical single-cell nano- mechanical measurements. Moreover as expected from the literature data [29,34], YM measured with sharp pyramidal tips is typically higher than the one measured using colloidal probes. This difference is clearly seen in the case of lung spheroids.

Based on the distribution of FCs collected within each measured spheroid, we did not observe a clear bimodal distribution of data, which could suggest the presence of elements with different rigidity (**Supplementary Figure S1**). Therefore, we can state that no influence of the presence of ECM or cell nucleus inside the specific cell type was detected in the mechanical properties of these spheroids. However, the spheroids formed from healthy fibroblasts (NHLF cells) were significantly stiffer than those from lung cancer (A549 cells). The corresponding YM was 7.60 ± 1.35 (n = 30 maps) versus 3.30 ± 0.36 (n = 36 maps) and 6.90 ± 0.58 (n = 30 maps) versus 3.70 ± 0.51 (n = 29 maps) for low and high indentations. The same observation regarding the independence of the measured YM value on the indentation depth was also made for the spheroids obtained from two colon cell lines (**Supplementary Figure S2**). Additionally, we did not observe any significant difference in rigidity between HT-29 and CCD18Co spheroids in both indentation ranges. Here, we measured 0.81 ± 0.07 kPa (n = 50 maps) for HT-29 spheroids and 0.76 ± 0.06 kPa (n = 51 maps) for CCD-18Co spheroids in low indentation range; in large indentation scale, the YM values were 0.90 ± 0.07 kPa (n = 50 maps) and 1.05 ± 0.08 kPa (n = 51 maps) for HT-29 and CCD18-Co spheroids, respectively. Both results indicate that in the case of colon cells, there is no difference in elastic properties between spheroids derived from cancerous cells and fibroblasts, which could be related to the absence of difference between their actin cytoskeleton organisations.

### 3.5. Tipless cantilevers

Finally, the elastic modulus of the spheroids was measured using a tipless cantilever (**Figure 3D**). The great advantage of tipless cantilevers is the large contact area due to the width of the cantilever (up to 100 µm) and its flat surface. Tipless cantilevers allow the indent of a spheroid as a whole rather than locally at the single- or few-cell level.

Using tipless cantilevers for the measurements on spheroids is an interesting approach; however, it raises questions regarding the contact mechanics and mechanical model that should be applied. To verify which model is most suitable, data from three spheroids were analysed to see which approach would be more suitable for further analysis (**Supplementary Figure S3**). Three different models and assumptions were used: the Hertz model for CPs with R equal to the radius of the spheroid, the Hertz model for CPs with R equal to half of the width of the cantilever, and the Hertz model cylindrical probe with R equal to half of the width of the cantilever.

First, we used the standard Hertz model but used an average radius of spheroids taken for measurements that day. This is equivalent to considering a deformable sphere (the spheroid) pushed against a rigid flat surface (the tipless cantilever, **Figure 1**). Second, we have used the standard Hertz model but assuming that the cantilever can be treated as a sphere with an effective diameter equal to the width of the cantilever. The third option was to use the flat punch model, as discussed above, with half of the cantilever width taken as radius a/2≍50 µm. However, the tipless cantilever does not provide a constant contact area, like in the case of truly cylindrical tips. The contact region can be considered an ellipse, with one axis determined by cantilever width and the other axis expected to increase with indentation. As a result, obtained data analysed using the flat punch approximation provided lower values of Young’s modulus than other models (**Supplementary Figure S3**). Giannetti et al. 2020 performed similar research on spheroids made of T24 (transitional cell carcinoma). The obtained elastic modulus was in the range of 100-500 Pa. Elastic modulus measured in this research (using the Hertz model with the radius of the spheroid, HertzSPH) were quite similar: 158.1 kPa for CCD-18Co and 213.2 kPa for HT-29 spheroids, while results obtained using the cylindrical model were lower than reported in Giannetti et al. 2020 and for Hertz model used in these studies [21]. This confirms that using the model for a cylindrical tip is not optimal while indenting with a tipless cantilever. Considering the results of data analysis, the most suitable and reliable model for further studies is the Hertz model for CPs, with a radius equal to the radius of spheroids. Both quantitative values of YM and consideration of contact mechanics favor this approach. The most important observation is that the values of YM are much smaller than modulus values when pyramidal and spherical probes were considered (**Figure 3**). The moduli range was 0.08 ± 0.006 kPa (n = 40 maps) and 0.25 ± 0.04 kPa (n = 30 maps) for spheroids formed respectively by A549 cells and NHLF fibroblasts. Comparison between the two colon cell lines revealed no differences, as mean values were 0.14 ± 0.01 kPa (n = 51) for HT-29 spheroids and 0.16 ± 0.01 kPa (n = 51) for CCD-18Co spheroids.

As described in the method section, we analysed the data for the tipless cantilevers by using the Hertz model for the sphere-plane contact geometry using the average radius of the spheroids (approximately 150 μm) as the radius R in Equation 1 and considering the tipless cantilever as a rigid flat surface. The other methods tested to simulate an effective sphere on flat geometry did provide similar results (see **Supplementary Figure S3**), nevertheless since these methods are based on rather crude geometrical assumptions we decided not to consider those results in the discussion. Overall, our results show that the lung spheroids formed by NHLF cells have a higher value of YM than those formed by the cancerous A549 cells, independent of the type of AFM probes. However, the magnitude of changes was related to the probing geometry and the most significant difference was observed for spherical probes at large 4-5 microns indentations. On the other hand, AFM measurements with different probes did not reveal any differences between colon cell lines studied here (**Figure 3** and Supplementary Figure S4).

The analysis of tumour-derived spheroids of A549 and HT-29 cell lines revealed that the outcome of the measurements could be different at different scales. The outcome of measurements with the MLCT tip was that A549-derived spheroids are stiffer than HT-29 spheroids, while measurements with a tipless cantilever showed the opposite results (**Supplementary Figure S5 A-C**). In the case of spheroids made of the fibroblast, NHLF spheroids were significantly stiffer, however, in measurements with tipless cantilever, these differences were more pronounced (**Supplementary Figure 5 D-F**).

## 4. Conclusions

Using various geometries of AFM tips allowed us to study the mechanical properties of spheroids at different length scales, from within the single cell to the multi-cellular level. It is known that the measured YM may depend on tip geometry, which is related to the contact area and the generated stress [34,40]. We observed indeed that the measured YM values were higher for sharper (pyramidal) tips in the case of all 4 types of spheroids. However, we observed the same relative trend among the different spheroids, irrespective of the probe used, especially when a larger and more regular contact area was used, i.e., for CPs and tipless cantilever.

It was observed before that cells in 3D can have different rigidity depending on their location. In the work of Vyas et. al. the YM of cells on the surface of the spheroid was higher than in the inner part of the proliferation zone [41]. Our results highlight the importance of selecting the right probe for studying the mechanical characteristics of biological samples, knowing that each component has a different YM value. The right choice of the AFM probe will depend on what information we want to extract: the average value of YM that includes the contribution of all sample elements or if we want to measure the rigidity of a single element within the sample. When measuring large and mechanically heterogeneous samples, the tip shape of the AFM probes influences the estimation of Young’s moduli; tips providing larger and regular contact areas should be preferred.

It is well known that cells can sense the rigidity of the microenvironment, which affects their morphology and behaviour [42–44]. This is why using spheroids as a study model is more advantageous than standard 2D culture. The cell’s response to the rigidity of the surrounding medium may vary between different cell types [42]. This might be linked to the organisation of the actin in the cells. We observed the difference in the organisation of actin fibers between spheroids of A549 and NHLF cell lines, and later we measured different values of YM. The differences in the rigidity were observed in the case of all measurements, using different geometries of the probes. By comparing the HT-29 and CCD-18Co spheroids, we observed no difference in the actin organisation. The cells in CCD-18Co spheroids showed similar morphology to cells in HT-29. This aligns with the results of AFM nanoindentation measurements, where no differences in rigidity were observed. Comparing lung fibroblasts with colon fibroblasts showed that the first ones are stiffer (**Figure S5 D-F**), which correlates with the information that a well-organised actin cytoskeleton increases the elastic modulus [45].

By comparing results obtained using different tip geometries and contact mechanics models, it can be seen that with the increase of radius used in the contact mechanics model, the obtained YM is significantly lower. Larger tips denote larger contact area and interaction volume, thus, the measured area is more averaging. Thus, it was possible to study the elastic properties of spheroids on a multi-cellular level while using pyramidal tips allowed us to extract values corresponding to individual elements of the system (i.e., single cells) in the spheroid outer layers. This comparison revealed that using different contact geometries in the fitting procedure results in significantly different YM values (**Supplementary Figure S4**), highlighting the importance of a precise experiment design and choice of the AFM probe for measurements.

Since the nature of the cells is linked to their internal structure, i.e., to all components constituting the cell, some of them will have a minor effect on the overall mechanics of the cell, and some will be dominant The overall conclusion is that regardless of the cantilever type, the most rigid spheroids were formed from lung fibroblasts, whose cytoskeleton was characterized by F-actin filaments located over a whole cell body. Spheroids composed of rounded cells with actin filaments located at their periphery were more compliant. Moreover, the choice of cantilever type is strongly bound to the relative difference between spheroid types. The difference between the mechanics of lung spheroids formed by spindle-like and round cells was the largest for cantilevers with small CPs, while an analogous large difference for spherical cells forming both lung and colon spheroids was observed when cantilevers with pyramidal tips were used. Using tipless or cantilevers with large CPs flattens the difference between different spheroid types. Our results suggest the dominant role of the actin cytoskeleton because, in our measurements, we are detecting the cells from the outer proliferative layer and not the space between them. For this reason, even if we probe at the large indentation, the response will still be dominant from actin filaments.

## Acknowledgement

This research was funded by the European Union’s Horizon 2020 research and innovation program under the Marie Skłodowska-Curie Action grant agreement No. 812772, project Phys2Biomed

## Supplementary

### SUPPLEMENTARY GRAPHS

**Supplementary Figure 1:**
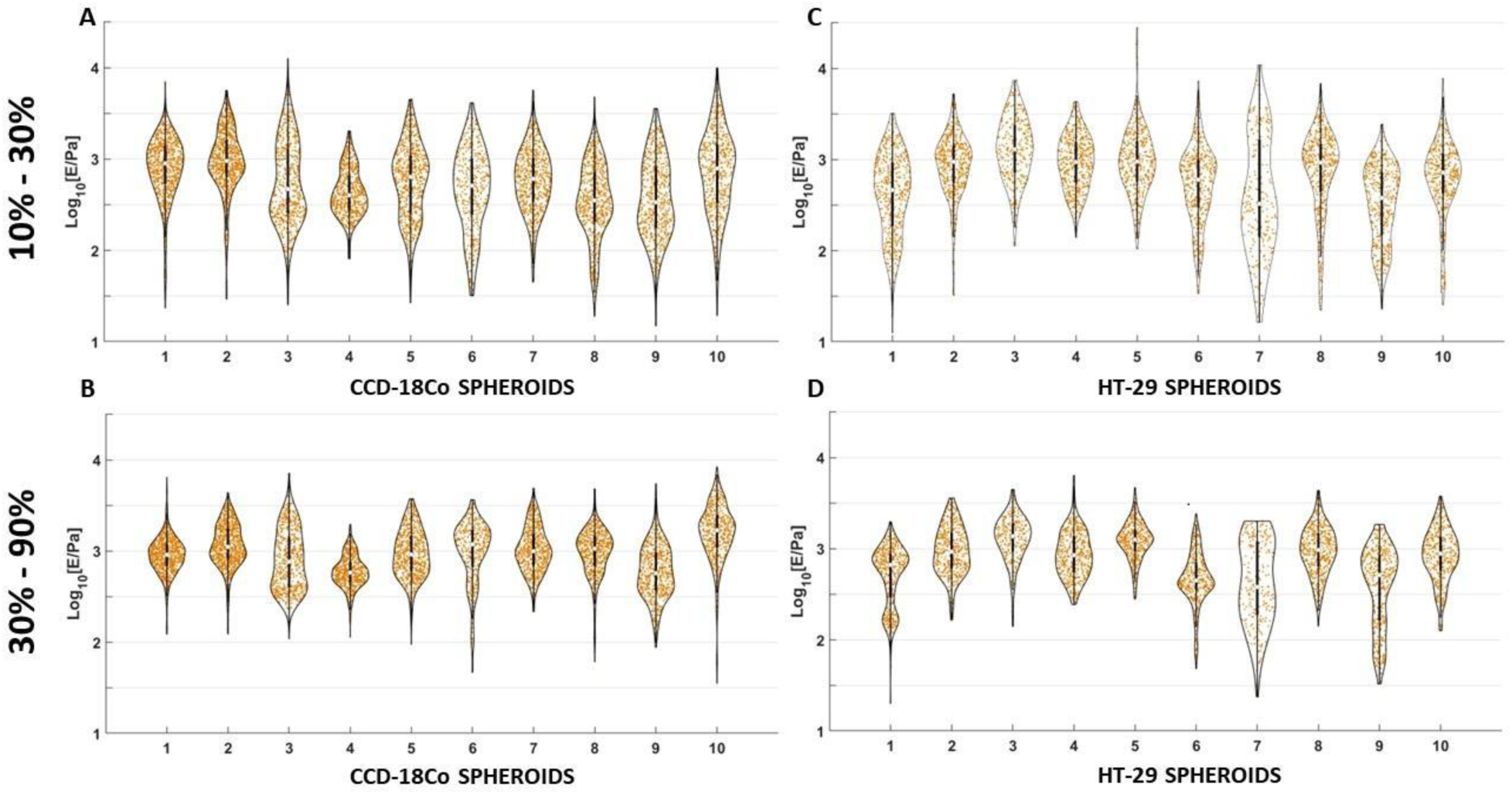
Violin plots presenting median YM values of each spheroid and distribution of YM obtained from single FCs. CCD-18Co measured with indentation of 10%-30% (A); CCD-18Co measured with indentation of 30%-90% (B); HT-29 measured with indentation of 10%-30% (C); HT-29 measured with indentation of 30%-90% (D)

**Supplementary Figure 2:**
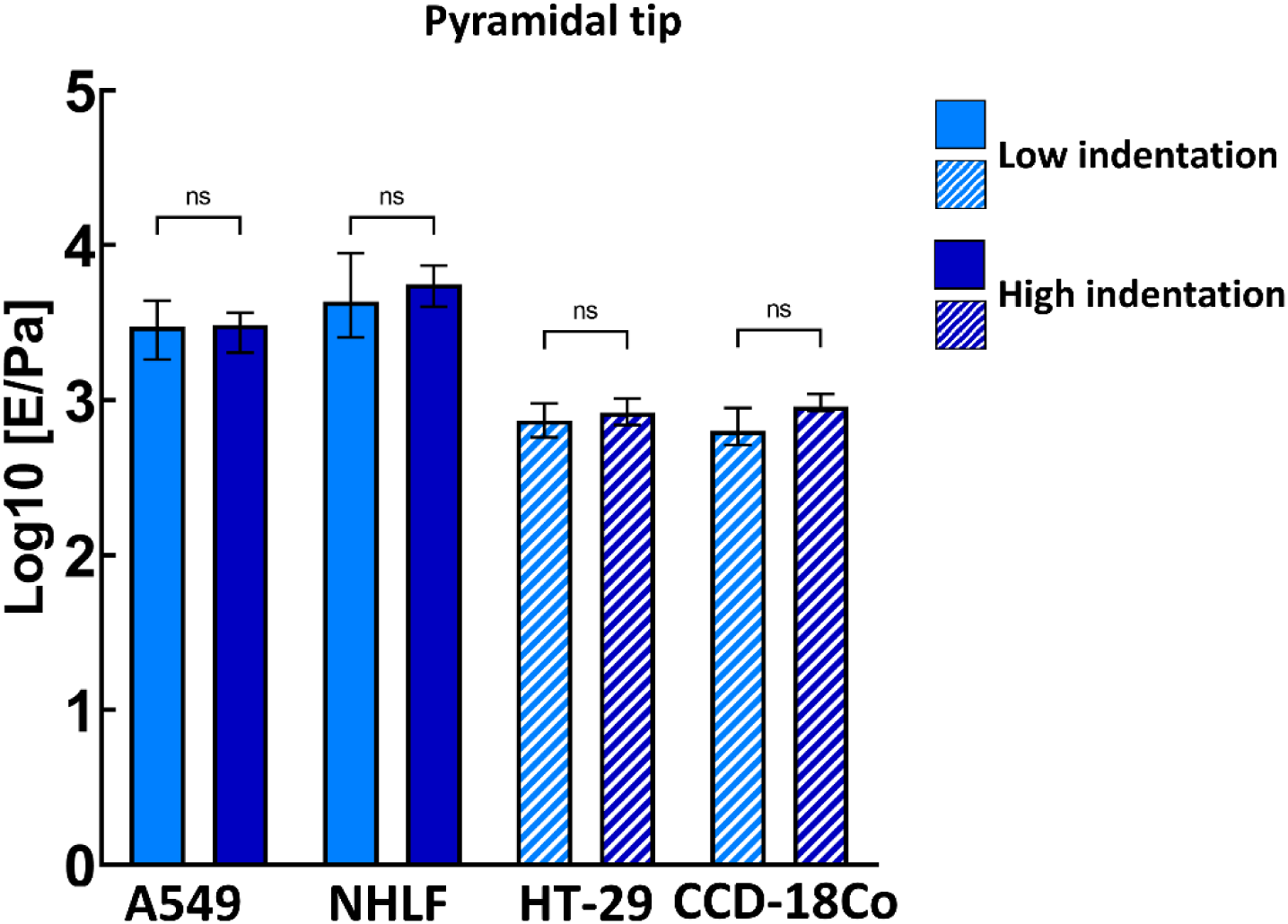
Comparison of two indentation ranges applied in measurements with pyramidal tip (**** p < 0.0001, ns - not statistically significant).

**Supplementary Figure 3:**
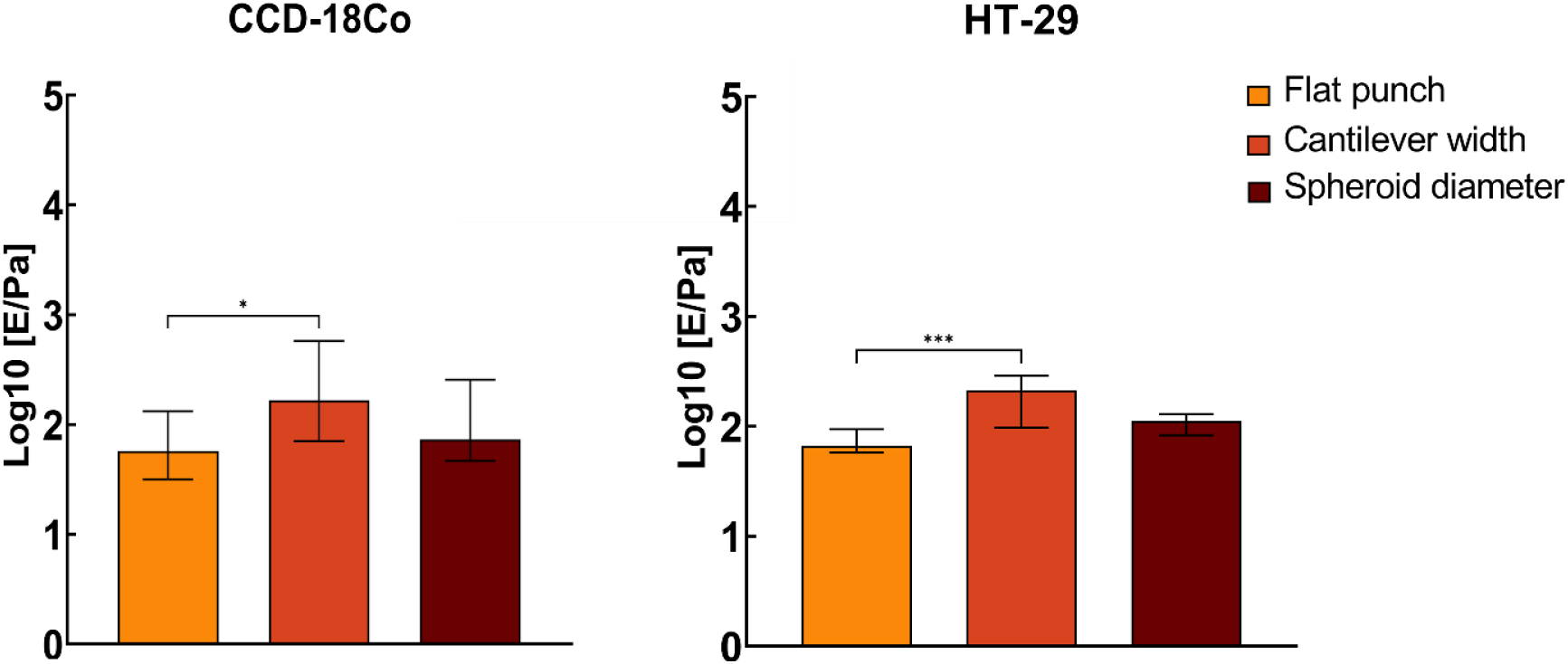
Results of measurements with tipless cantilever analysed using three different approaches in contact mechanics (**** p < 0.0001, ns - not statistically significant).

**Supplementary Figure 4:**
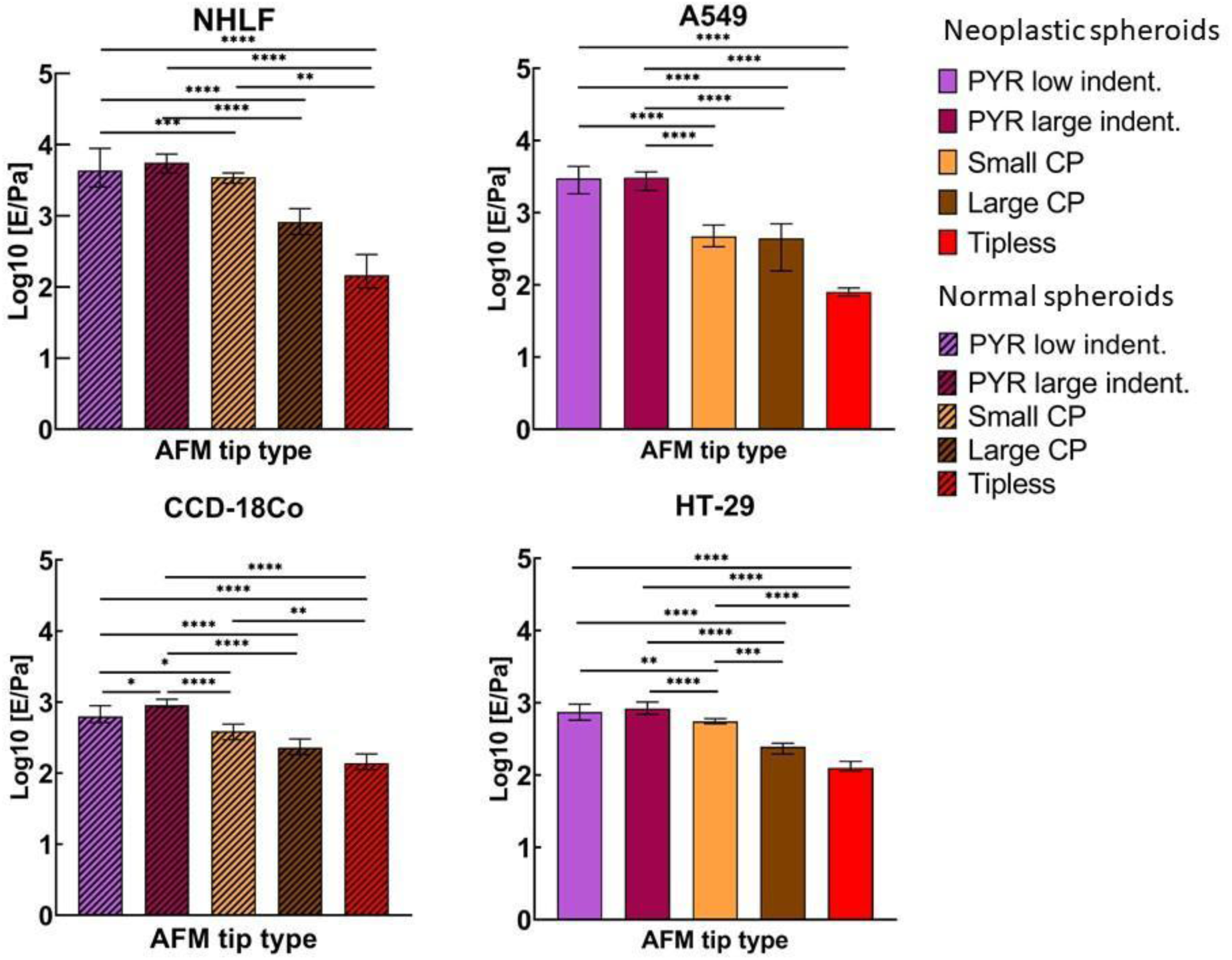
The comparison between YM median values measured with different AFM tip geometries within each spheroid type. The graphs show a median with a 95% CI of log values (median of the medians). Ordinary ANOVA test with Turkey’s multiple comparison test was performed to verify the statistical significance of differences.

**Supplementary Figure 5:**
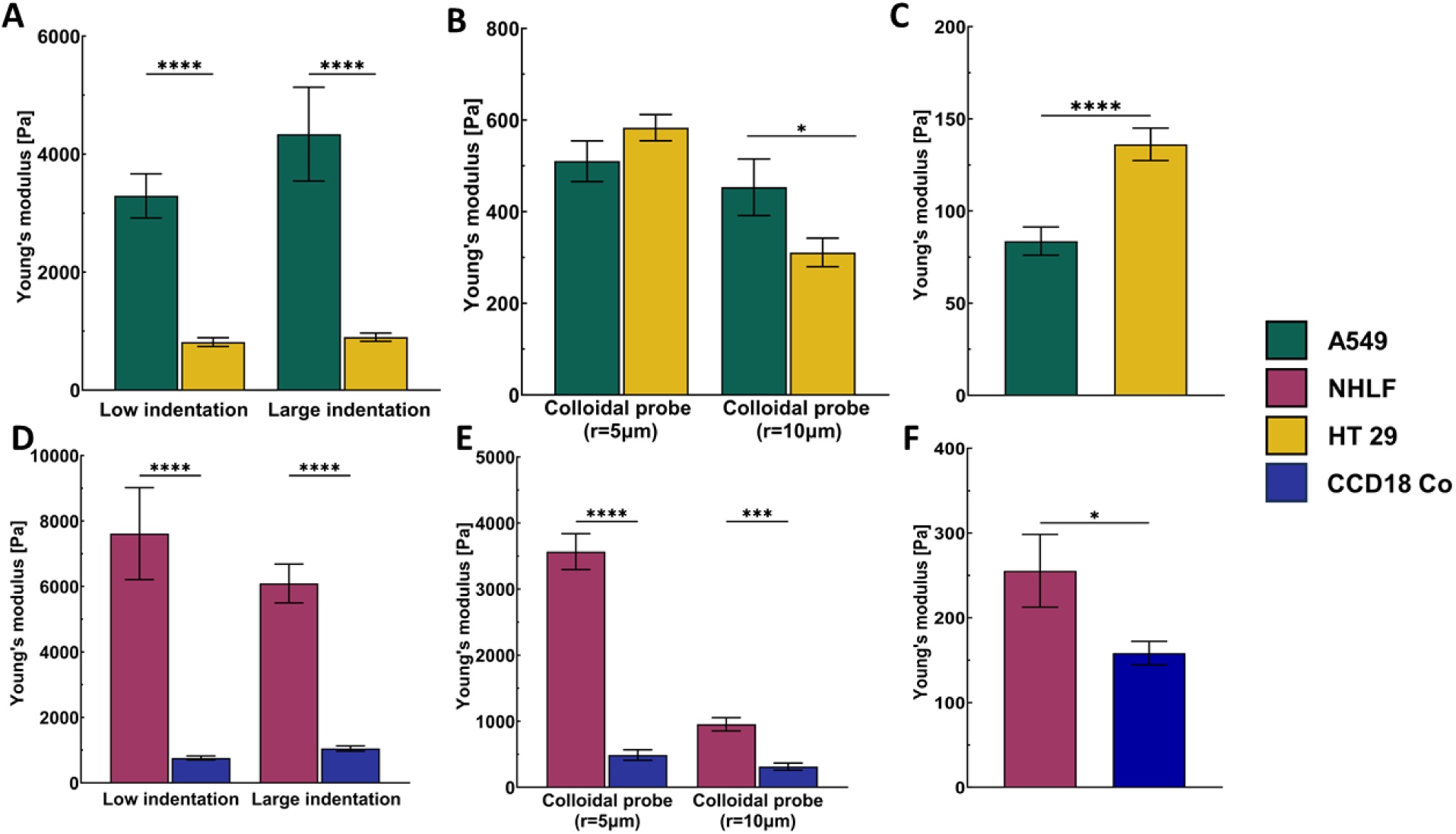
Comparison of YM values between cancer spheroids (A-C) and between fibroblast-derived spheroids (D-F). The measurements were done with MLCT cantilevers at low and large indentations (A,D), spherical probes (B,E), and tipless cantilevers (C,F). Each bar represents the mean value ± standard error of the mean. (**** p < 0.0001, ns - not statistically significant).

## Notes

### Competing Interest Statement

The authors have declared no competing interest.

